# Extremely low level of genetic diversity in *Gentiana yakushimensis*, an endangered species in Yakushima Island, Japan

**DOI:** 10.1101/2022.05.18.492409

**Authors:** Naohiro I. Ishii, Shun K. Hirota, Yoshihiro Tsunamoto, Ayumi Matsuo, Harue Abe, Yoshihisa Suyama

**Author notes:** **Correspondence to:** Naohiro I. Ishii, Graduate School of Environment and Information Sciences, Yokohama National University, 79-7 Tokiwadai, Hodogaya, Yokohama, Kanagawa 240-8501, Japan.

## Abstract

Rare or endangered species largely contributing to global biodiversity are essential components of ecosystems. Because genetic diversity depends not only on apparent population size but also demographic processes, the conservation priority of endangered species based on population genetics needs to be assessed by comparison with common congeners and demographic events. In this study, we performed population genetic analysis for *Gentiana yakushimensis* Makino, a rare and endangered plant species with an isolated distribution on Yakushima Island, Japan. We performed comparison of genetic diversity with the congener (*Gentiana triflora* Pall. var. *japonica* (Kusn.) H.Hara), demographic inference, and bottleneck test based on genome-wide single nucleotide polymorphisms. The result was that *G. yakushimensis* had an extremely low genetic diversity level and a high inbreeding coefficient level compared to those of the congener. The population of *G. yakushimensis* was estimated to experience an expansion during the Last Glacial Maximum and a recent bottleneck by the demographic inference and the bottleneck test. Therefore, the low level of genetic diversity of this species might have resulted from the impact of the recent bottleneck and the long-term maintenance of a small population size. Due to the long lifespan of the genus *Gentiana*, the current level of genetic diversity probably does not reflect a large part of recent demographic events, which requires long-term monitoring of changes in genetic diversity to identify the time lag between the reduction in apparent population size and genetic diversity as a future outlook.

## Introduction

Rare or endangered species, largely contributing to global biodiversity, are essential components of various ecosystems (Enquist et al., 2019). Most previous studies have demonstrated that rare species with limited geographic distributions tend to have lower levels of genetic diversity than common species because of their small effective population size (Broadhurst et al., 2017; Chung et al., 2018; Gibson, Rice, & Stucke, 2008; Hamrick & Godt, 1996; Nybom & Bartish, 2000). The low level of genetic diversity in a species or a population can trigger its vulnerability because it could generally result in the reduction in adaptive potential to environmental changes and resilience or resistance to environmental extremes (Reusch, Ehlers, Hammerli, & Worm, 2005; Willi, Van Buskirk, & Hoffmann, 2006). Inbreeding depression associated with low genetic diversity can also reduce reproductive fitness (Losdat, Chang, & Reid, 2014; O’Grady et al., 2006; Reed & Frankham, 2003). Thus, it is reasonable to globally evaluate the conservation priority of rare or endangered species based on apparent population size and genetic diversity (Garner, Hoban, & Luikart, 2020). Accumulation of genetic information for thousands of endangered species can lay the foundation for biodiversity conservation and conservation prioritization.

The genetic diversity of rare species is not always low, and it can be species-specific because it depends on apparent population size and demographic processes or life history traits (Ellegren & Galtier, 2016; Hamrick & Godt, 1996; Nybom & Bartish, 2000; Thiel-Egenter et al., 2009). For example, previous studies have demonstrated that species with a long lifespan tend to have higher levels of genetic diversity than those with short lifespans (Kort et al., 2021; Reisch & Rosbakh, 2020). Past and current demographic events, such as population bottlenecks or expansion due to environmental and anthropogenic fluctuations, can affect the current genetic diversity while comparing closely related species with similar life history traits (Ellegren & Galtier, 2016). For example, the species widely distributed in the past can become rare or endangered due to disturbances, resulting in the low level of genetic diversity (e.g. Li et al., 2014). Conversely, in isolated habitats such as islands, a small population size can be maintained for long term due to limited immigrants, which was estimated to be a factor of low genetic diversity (Gonzalez, Cerón-Souza, Mateo, & Zardoya, 2014). Therefore, estimating demographic history allows us to examine which current level of genetic diversity is determined by historical or recent events.

Alpine plants, including many rare or endangered species, are the targets for which conservation prioritization based on genetic aspects is required (Noroozi et al., 2018; Perrigo, Hoorn, & Antonelli, 2020), as these plants are vulnerable to climate change and anthropogenic disturbances (Guisan & Theurillat, 2000; Thuiller, Lavorel, Araújo, Sykes, & Prentice, 2005). Previous studies have demonstrated that the current levels of genetic diversity of alpine plants are strongly affected by several demographic events, such as expansion or reduction of distribution range after the Last Glacial Maximum (LGM; 21,000 years ago) (Loïc Pellissier et al., 2016), geographical isolation among populations by the distribution patterns restricted to high-altitude areas (Reisch & Rosbakh, 2020), and direct human activities such as trampling by tourists and illegal digging (Banks et al., 2013; Feng, Liu, Chiang, & Gong, 2017). Therefore, inferring historical demography allows us to examine the influence of historical demographic events on current genetic diversity levels and accurately assess the conservation status of rare alpine species.

*Gentiana yakushimensis* Makino (Figure 1) is an endangered plant species in Japan (Japanese Ministry of Environment, 2020) and endemic to Yakushima Island (Iwatsuki, Yamazaki, Boufford, & Ohba, 1993). On Yakushima Island, designated a UNESCO World Heritage Site, this species grows only in limited alpine zones covered by granite rocks with low vegetation cover in the central mountains (Figure 2). Illegal digging for ornamental horticulture has drastically reduced its population (Japanese Ministry of Environment, 2020). *G. yakushimensis*, or its ancestor, was estimated to colonize Japan 4–1 million years ago with the Qinghai–Tibet Plateau as the origin (Appendix S3 in Favre et al., 2016). The taxonomic position of *G. yakushimensis* is discussed. Recently, phylogenetic analysis based on chloroplast DNA regions suggested that *G. yakushimensis* is close to the section *Pneumonanthe* (Mishiba et al., 2009). Moreover, phylogenetic analysis covering all *Gentiana* sections supported the inclusion of *G. yakushimensis* in *the Pneumonanthe*/*Cruciata* lineage (Favre et al., 2016). The previous phylogeographic studies of alpine species have demonstrated range shifts or persistence during and after the LGM not only in Japan (e.g Ikeda, Higashi, Yakubov, Barkalov, & Setoguchi, 2014; Ikeda, Yakubov, Barkalov, Sato, & Fujii, 2020), but also globally (e.g Allen, Marr, Mccormick, & Hebda, 2015; Ikeda et al., 2017; Schönswetter, Stehlik, Holderegger, & Tribsch, 2005). Thus, by estimating the population demography of *G. yakushimensis*, which distributed in the subalpine region, we can examine whether the current level of genetic diversity resulted from recent and/or historical demographic events.

**FIGURE 1.**
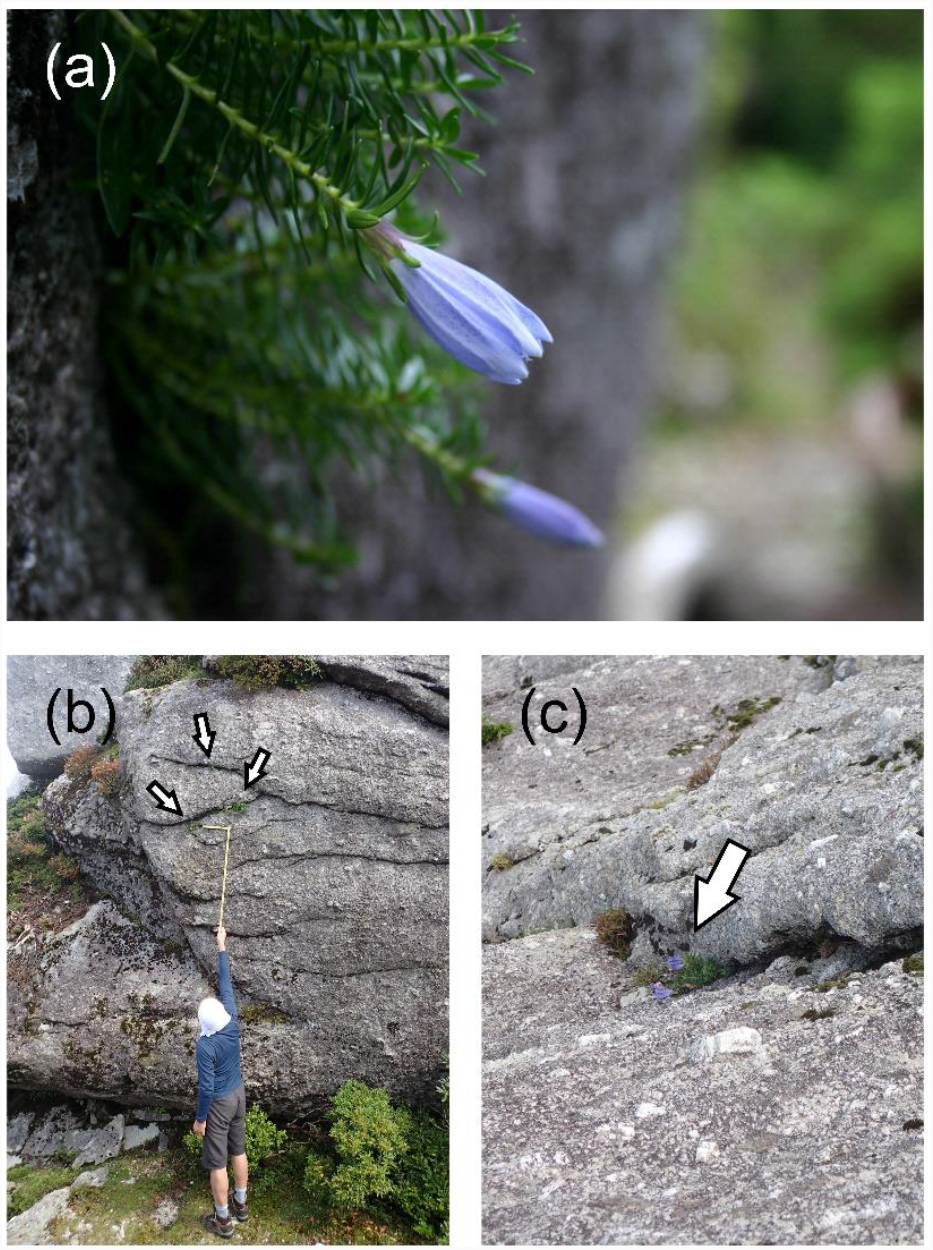
Photograph of (a) *Gentiana yakushimensis* and (b-c) its habitat. The white arrows in the panels (b) and (c) point to *Gentiana yakushimensis*.

**FIGURE 2.**
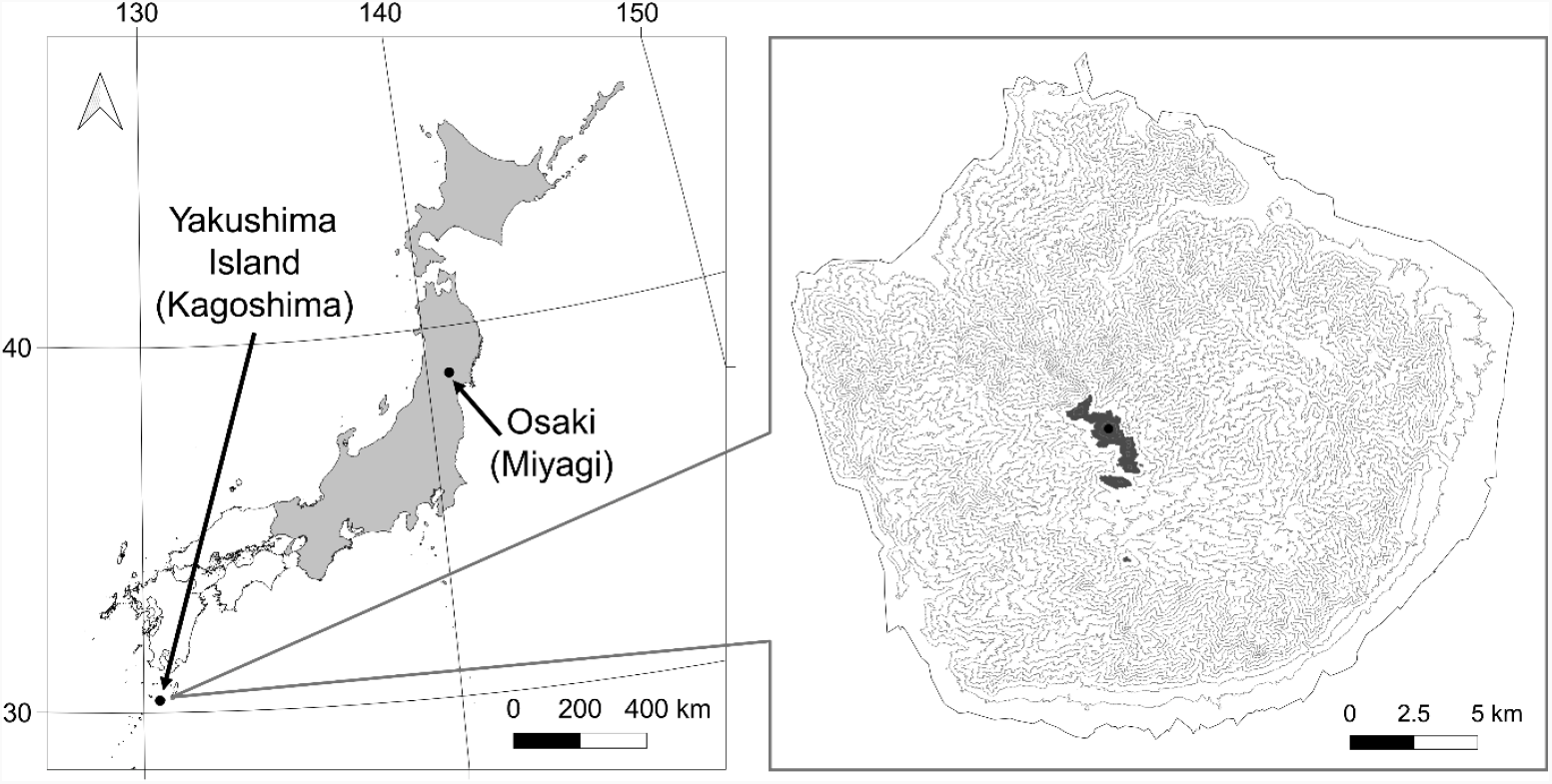
Map of Yakushima Island and the distribution area of *Gentiana yakushimensis*. The gray part in the right map denotes the area above 1,700 m above sea level. In the left map, the gray part denotes the distribution area of *G. triflora*. The contour interval is 50 m.

Here, we assessed the genetic diversity and demographic history of *G. yakushimensis* based on genome-wide single nucleotide polymorphisms (SNPs) to identify their conservation status. We performed 1) the comparison of genetic diversity with a congener (*Gentiana triflora* Pall. var. *japonica* (Kusn.) H.Hara), 2) the evaluation of genetic differentiation across Yakushima Island, and 3) the demographic inference of *G. yakushimensis* to identify whether the determinant of the current level of genetic diversity is recent or historical, aiming to provide the genetic information of the endangered species, supporting decision-making in conservation implications, and prioritizing.

## Materials and Methods

### Studied species

*G. yakushimensis* (Gentianaceae) is a perennial herb endemic to Yakushima Island, Japan (Chen & Wang, 1999; Ho & Liu, 2001; Iwatsuki et al., 1993). Although Ho (1988) recorded the occurrence of *G. yakushimensis* in Taiwan, China based on a single specimen (PE No. 81854; https://www.cvh.ac.cn/spms/detail.php?id=ec90580f), Chen and Wang (1999) denied the distribution of this species in Taiwan because the specimen lacks collection labels and no other specimens from Taiwan exist. *G. yakushimensis* has a minimal and patchy distribution; it frequently appears on granite rocks around the tops of the central mountains, at altitudes ranging from 1,700 to 1,900 m (Figure 2). *G. yakushimensis* was pollinated by bees, which were likely to be Andrenidae and Halictidae (H. Abe, personal observation), and its seeds with no wings were assumed to be dispersed by gravity and wind. The population size of *G. yakushimensis* has decreased due to illegal digging in recent years (Japanese Ministry of the Environment, 2014). In addition, feeding damage by sika deer (*Cervus nippon yakushimae*) might also be a factor in population size reduction. In Yakushima Island, grazing by sika deer has affected vegetation from lowlands to subalpine zones (Takatsuki, 1990; Tsujino & Yumoto, 2004). Takatsuki (1990) demonstrated that sika deer relied on forbs for about 5 % of their total diet in the subalpine area of Yakushima Island, indicating that *G. yakushimensis* could be affected by grazing. Thus, *G. yakushimensis* has been listed in the Japanese Red List as Endangered (EN) (Japanese Ministry of Environment, 2020) and conserved under the “Act on Conservation of Endangered Species of Wild Fauna and Flora” since 2016.

*Gentiana triflora* var. *japonica* was used to compare the genetic diversity of *G. yakushimensis. G. triflora* var. *japonica* is a relatively common species widely distributed in the mountainous area from Hokkaido to the Kinki region of Japan (Figure 2). This species is one of the most closely related to *G. yakushimensis* based on the chloroplast DNA region (Favre et al., 2016; Mishiba et al., 2009).

We collected leaf samples from all individuals of *G. yakushimensis* (560 individuals), which had grown in accessible habitats, for DNA analysis at Yakushima Island in August from 2009 to 2010. Small individuals or individuals with few leaves were not collected to conserve *G. yakushimensis*. When multiple individuals grew at the same coordinates, one individual was selected for DNA analysis. Leaf samples were collected with permission from the prefecture and ministries. The leaf samples of *G. triflora* were collected from nine individuals as congeners from an area of approximately 100 m^2^ in Osaki, Miyagi Prefecture, in the northern area of Japan (Figure 2).

### DNA extraction, amplification, and sequencing

Total genomic DNA was extracted from frozen leaf samples using the modified cetyltrimethylammonium bromide (CTAB) method (Doyle, 1991). For DNA analysis, we used 353 and 9 samples of *G. yakushimensis* and *G. triflora*, respectively (Table 1). The samples of *G. yakushimensis* for the analysis were at least 3 m away from each other because whether the different shoots in granite crevices were from the same individual could not be clearly distinguished.

**TABLE 1.** Indices of genetic diversity and inbreeding coefficient parameters of *Gentiana yakushimensis* and *G. triflora*. r, the parameter of the minimum sharing rate among individuals in the ‘populations’ command of Stacks v2.52; π, nucleotide diversity; *H*_O_, observed heterozygosity; *H*_E_, expected heterozygosity; *F*_IS_, inbreeding coefficient. *P*-values were based on pairwise Wilcoxon rank sum test. *P*_0_ denotes *P*-value of one-sample Wilcoxon rank sum test whether *F*_IS_ was significantly different from zero. *, *P* < 0.05; **, *P* < 0.01; ***, *P* < 0.001.

| Common/<br>all<br>loci | Species | No. of<br>samples | $r$ | No. of loci | No. of<br>polymorphic<br>loci | No. of<br>Sites | No. of<br>polymorphic<br>sites | $\pi$ | | | $H_O$ | | | $H_E$ | | | $F_{IS}$ | | | |
| --- | --- | --- | --- | --- | --- | --- | --- | --- | --- | --- | --- | --- | --- | --- | --- | --- | --- | --- | --- | --- |
| | | | | | | | | Mean | SE | $P$ | Mean | SE | $P$ | Mean | SE | $P$ | Mean | SE | $P_0$ | $P$ |
| Common | *G. yakushimensis* | 353 | 0.5 | 89 | 47 | 7,237 | 72 | 0.00020 | 0.00008 | *** | 0.013 | 0.005 | *** | 0.020 | 0.008 | *** | 0.222 | 0.045 | *** |  |
| Common | *G. triflora* | 9 |  | 89 | 47 | 7,237 | 72 | 0.00135 | 0.00030 |  | 0.121 | 0.030 |  | 0.125 | 0.023 |  | 0.048 | 0.076 |  |  |
| Common | *G. yakushimensis* | 353 | 0.6 | 69 | 35 | 5,623 | 51 | 0.00009 | 0.00002 | ** | 0.006 | 0.002 | ** | 0.009 | 0.002 | *** | 0.263 | 0.054 | *** | * |
| Common | *G. triflora* | 9 |  | 69 | 35 | 5,623 | 51 | 0.00103 | 0.00028 |  | 0.100 | 0.030 |  | 0.105 | 0.025 |  | 0.058 | 0.075 |  |  |
| Common | *G. yakushimensis* | 353 | 0.7 | 51 | 25 | 4,150 | 34 | 0.00005 | 0.00002 | ** | 0.004 | 0.001 | * | 0.006 | 0.002 | ** | 0.274 | 0.069 | *** | * |
| Common | *G. triflora* | 9 |  | 51 | 25 | 4,150 | 34 | 0.00095 | 0.00031 |  | 0.113 | 0.041 |  | 0.108 | 0.031 |  | -0.004 | 0.098 |  |  |
| All | *G. yakushimensis* | 353 | 0.5 | 2,124 | 487 | 172,094 | 784 | 0.00055 | 0.00003 | *** | 0.084 | 0.004 | *** | 0.121 | 0.005 | *** | 0.341 | 0.014 | *** | *** |
| All | *G. triflora* | 9 |  | 1,475 | 302 | 116,730 | 401 | 0.00162 | 0.00008 |  | 0.329 | 0.009 |  | 0.436 | 0.002 |  | 0.230 | 0.022 | *** |  |
| All | *G. yakushimensis* | 353 | 0.6 | 1,758 | 387 | 142,510 | 562 | 0.00041 | 0.00003 | *** | 0.070 | 0.005 | *** | 0.103 | 0.005 | *** | 0.358 | 0.017 | *** | *** |
| All | *G. triflora* | 9 |  | 1,181 | 222 | 93,622 | 297 | 0.00147 | 0.00009 |  | 0.311 | 0.009 |  | 0.430 | 0.003 |  | 0.263 | 0.023 | *** |  |
| All | *G. yakushimensis* | 353 | 0.7 | 1,429 | 292 | 115,877 | 402 | 0.00033 | 0.00003 | *** | 0.058 | 0.005 | *** | 0.095 | 0.006 | *** | 0.387 | 0.019 | *** | *** |
| All | *G. triflora* | 9 |  | 945 | 155 | 74,977 | 202 | 0.00125 | 0.00009 |  | 0.315 | 0.011 |  | 0.435 | 0.004 |  | 0.264 | 0.027 | *** |  |

To evaluate the genetic diversity of *G. yakushimensis* and *G. triflora*, we performed *de novo* single-nucleotide polymorphism (SNP) detection using multiplexed inter-simple sequence repeat (ISSR) genotyping by sequencing (MIG-seq) (Suyama et al., 2021; Suyama & Matsuki, 2015). We prepared the MIG-seq library and sequenced it following the protocol described by Suyama et al. (2021). Briefly, ISSRs were amplified using primer set-1 (Suyama & Matsuki, 2015) in the first PCR, and then the index and adaptor sequences were added to both ends of the 1st PCR products in the second PCR. We sequenced 80 base pair-end reads using the Illumina MiSeq platform and MiSeq Reagent Kit v3 (150 cycles) (Illumina, San Diego, CA, USA). We skipped detection of the first 17 bases of Reads 1 and 2 using the “DarkCycle” option.

### Read assembling and SNP marker filtering

One base at each end of the processed reads was trimmed, and the reads were filtered based on the quality of reads and the adaptor sequences using Trimmomatic v.0.3.2 (Bolger, Lohse, & Usadel, 2014). Therefore, we performed the subsequent analysis using filtered reads with a length of 78 bp. Stacks v2.52 (Catchen, Amores, Hohenlohe, Cresko, & Postlethwait, 2011) which was used to process the filtered reads with the default parameter settings: the minimum number of identical reads required to create a stack (m = 3), the nucleotide mismatches between loci within a single individual (M = 2), and mismatches between loci when building the catalog (n = 1). The SNP genotype for each individual was exported with the following parameter setting using the ‘populations’ command in Stacks v2.52: the maximum observed heterozygosity, --max-obs-het = 0.6; minimum minor allele frequency, --min-maf = 2 / the number of individuals. For the dataset of both species (common loci) and individual species (common and species-specific loci; hereafter, referred to as “all loci”), the SNP genotype was exported by setting r (the parameter of the minimum sharing rate of SNP among individuals per population) at 0.5, 0.6, and 0.7, in addition to the above parameter settings to compare the genetic diversity of *G. yakushimensis* and *G. triflora*. We set up each species as a population to meet the r threshold for each species (p = 2 for the dataset of common loci and p = 1 for all loci in the ‘populations’ command). The comparison with the different values of the parameter r was performed because the number of SNPs of the common loci was low (less than 100) when the parameter r was 0.7. The estimation of genetic indices with low parameter r values leads to increase in missing data but ensures accuracy by increasing the number of SNPs (Hodel et al., 2017).

### Comparison of genetic diversity between G. yakushimensis and G. triflora

Genetic diversity was compared between *G. yakushimensis* and *G. triflora* for both the common and all loci. We calculated nucleotide diversity (π) based on fixed and variant sites using the ‘populations’ command of Stacks v2.52. The other indices, observed and expected heterozygosity (*H*_O_ and *H*_E_), and inbreeding coefficient (*F*_IS_), were determined using GenAlEx v6.51 (Peakall & Smouse, 2006). Wilcoxon rank sum tests were performed on these indices to test the significance of the difference between the species.

### Genetic differentiation of G. yakushimensis

Subsequent analyses were performed using the dataset of all *G. yakushimensis* loci. Kosman & Leonard’s (2005) pairwise genetic distance between *G. yakushimensis* individuals was calculated using the R package PopGenReport (Adamack & Gruber, 2014). Based on genetic and geographic distance matrices among individuals, we assessed spatial autocorrelation (isolation by distance, IBD) across the distribution area using the ‘mantel.correlog’ function in the R package vegan (Oksanen et al., 2020). To test for the presence of genetic structure, we performed principal coordinate analysis (PCoA) based on the genetic distance matrix using the ‘pcoa’ function in the R package ape (Paradis & Schliep, 2019).

### Inference of population demographic history

We inferred the population demographic history of *G. yakushimensis*. In the ‘populations’ command in Stacks v2.52, we exported SNP genotypes for the dataset of the all loci of *G. yakushimensis* with the change of the following parameter: minimum minor allele frequency, --min-maf = 0. Population demography was inferred by Stairway plot v2.1 (Liu & Fu, 2015), a model-flexible method of inferring historical changes in population size based on folded site frequency spectrums (SFS). SFSs were calculated using easySFS (https://github.com/isaacovercast/easySFS), and samples were projected downward to maximize the number of loci without missing data and number of individuals retained (Kreiner et al., 2019). The parameter settings were as follows: 67% of sites for training; 15 years per generation; 4.089 × 10^−8^ of mutation rate. We assumed that the lifespan (generation time) of *G. yakushimensis* is 15 years, according to which the lifespan was at least 15 years in a congener, *G. pneumonanthe* (Oostermeijer, Veer, & Nijs, 1994). The mutation rate was calculated by the relationship between genome size and mutation rate (Lynch, 2010), following Garot, Joët, Combes, Severac, & Lashermes (2019). The mean genome size of the genus *Gentiana* (3640.7 Mb) in the plant DNA C-values database (Pellicer & Leitch, 2020) used for this analysis.

Finally, the signatures of recent genetic bottleneck were assessed using the heterozygote excess test implemented in the software BOTTLENECK (Cornuet & Luikart, 1996; Piry, Luikart, & Cornuet, 1999) by INEST v2.2 (Chybicki & Burczyk, 2009). This method compares *H*_E_ in an empirical population to the heterozygosity that is expected in a population at mutation-drift equilibrium (*H*_Eq_). A transitory excess in *H*_E_ (measured as *ΔH* = *H*_E_ - *H*_Eq_) is assumed in recently bottlenecked populations. The significance of the heterozygote excess was evaluated using Wilcoxon signed-rank tests under an infinite alleles model (IAM) (Maruyama & Fuerst, 1985). Although IAM is not entirely appropriate for biallelic SNPs, this test was exploratorily conducted following recent studies (Camak, Osborne, & Turner, 2021; Kogura et al., 2011; Setzke, Wong, & Russello, 2022; Stojanova, Šurinová, Zeisek, Münzbergová, & Pánková, 2020).

## Results

The mean number of filtered reads per sample of *G. yakushimensis* and *G. triflora* was 92,960 and 58,900, respectively. The number of SNPs in the common loci between the two species was 72, 51, and 34, with r = 0.5, 0.6, and 0.7, respectively (Table 1). In the common and species-specific (all) loci of *G. yakushimensis*, the number of SNPs was 784, 562, and 402 with r = 0.5, 0.6, and 0.7, respectively (Table 1). On the other hand, the number of SNPs in all loci of *G. triflora* was 401, 297, and 202, with r = 0.5, 0.6, and 0.7, respectively (Table 1).

Compared to *G. triflora*, the genetic diversity indices (π, *H*_O_, and *H*_E_) of *G. yakushimensis* were significantly lower in all parameter settings (Table 1). When comparing *G. yakushimensis* with *G. triflora* for common loci with setting r = 0.7 in the ‘populations’ command, the former was 0.05 times for π and *H*_E_, and 0.03 times for *H*_O_ as much as the latter (Table 1). The inbreeding coefficients (*F*_IS_) of *G. yakushimensis* were significantly greater than zero for all parameter settings (Table 1). Also, it was significantly greater than those of *G. triflora* except for the common loci with setting r = 0.5 in the ‘populations’ command.

We observed a significant positive correlation between genetic and geographic distances among the individuals of *G. yakushimensis* (Mantel correlation coefficient, *r* = 0.25– 0.27, *P* < 0.001) (Figure 3a). However, an increase in genetic differentiation among the individuals was not detected when the pairwise geographical distance was over approximately 500 m. As a result of the Mantel correlogram, calculating Mantel correlation coefficients for each distance class, consistently positive correlations were shown up to approximately 140– 160 m (Figure 3b). PCoA did not demonstrate population genetic structure across the distribution area, although the first three axes of PCoA explained only 4–6 % of the whole genetic variation of *G. yakushimensis* (Figure 4, and figure S1).

**FIGURE 3.**
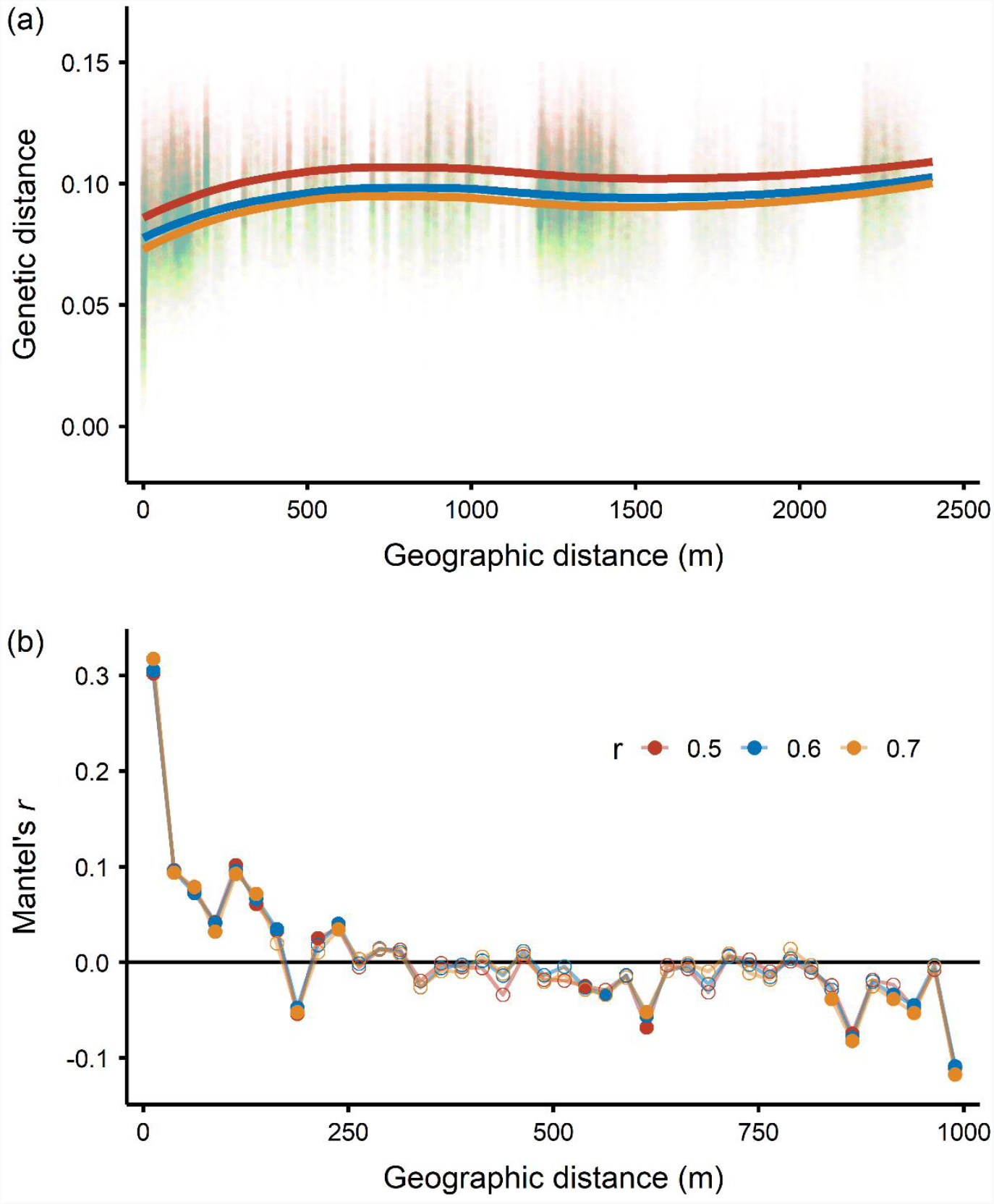
Relationship between pairwise genetic and geographic distance. (a) Relationship between individual-based pairwise genetic and geographic distance of *Gentiana yakushimensis* when the parameter r is 0.5, 0.6, and 0.7. (b) Mantel correlograms for the association between individual-based genetic and geographic distances. Filled symbols in the figure denote statistical significance.

**FIGURE 4.**
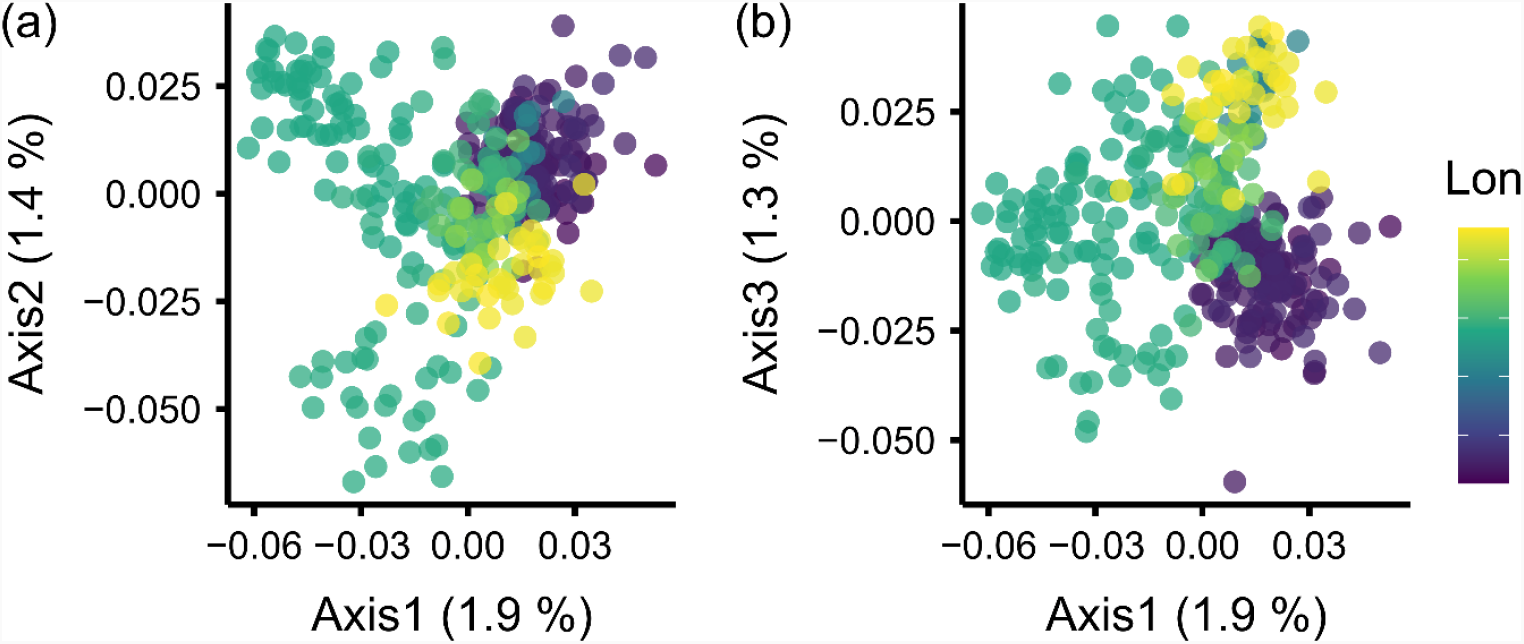
Principal coordinate analysis (PCoA) score plots based on individual-based genetic distance of *Gentiana yakushimensis* when the parameter r is 0.5. (a), axis 1-2; (b), axis 1-3. Colors denote the location (longitude) of each individual.

As a result of the demographic inference by the Stairway plot, the expansion of population size at 10,00040,000 years ago and the constant population size until recent years were estimated (Figure 5). The median of the population size of 5,0006,000 individuals was inferred to be maintained for at least 10,000 years. In addition, the 95% confidence intervals of population size were expanded downward from 1,700 to 10,000 at about 50 years ago (34 generations ago). These patterns were consistent among with the different setting of the parameter (r = 0.5, 0.6, and 0.7 in the ‘populations’ command in Stacks). As the results of the test of genetic bottleneck, the median of *ΔH* was significantly higher than zero (*H*_E_ was higher than *H*_Eq_) at each setting of the parameter (r = 0.5, 0.6, and 0.7), which exhibited that the population of *G. yakushimensis* showed significant heterozygote excess (Figure 6).

**FIGURE 5.**
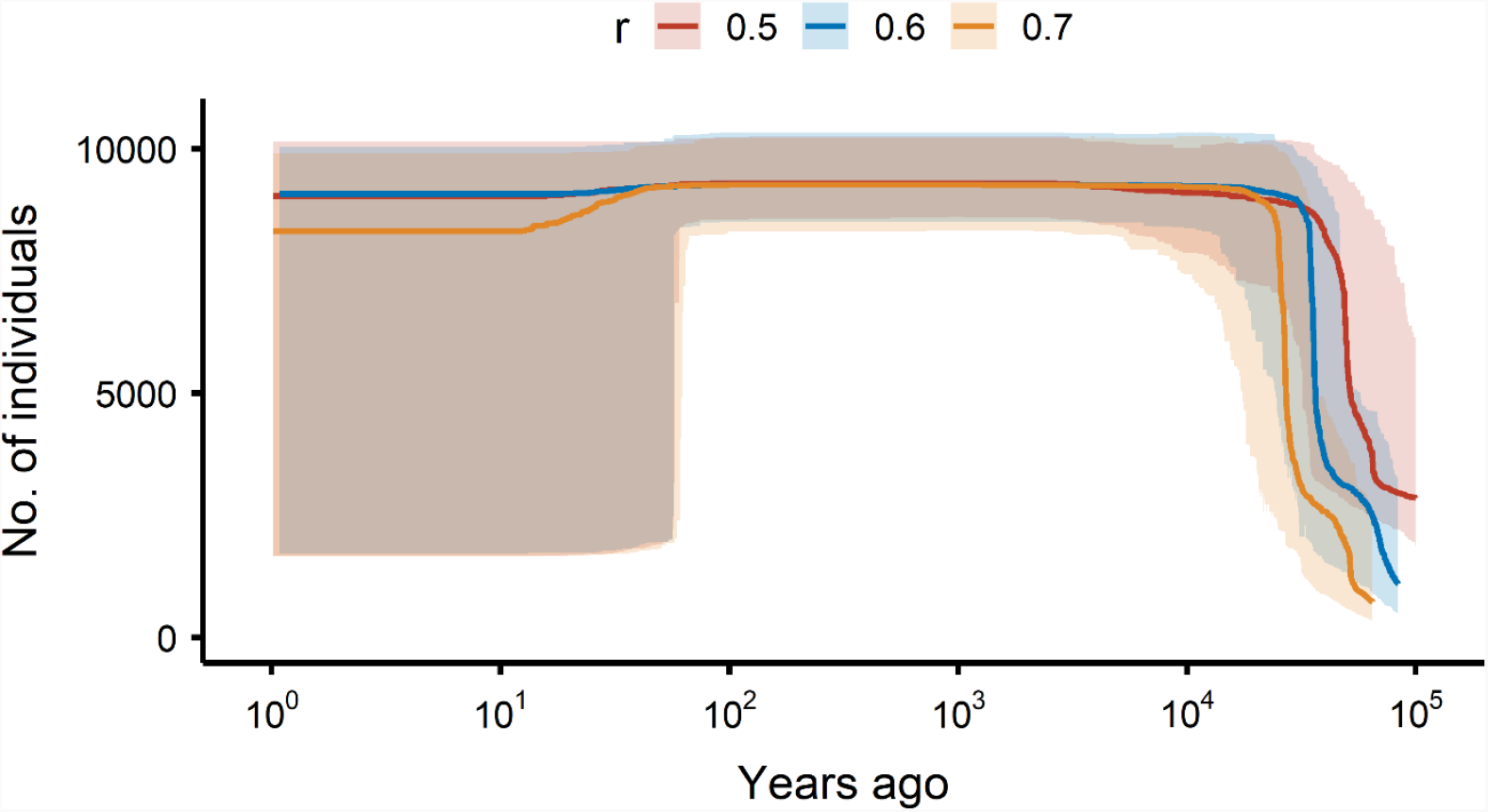
Demographic inference using stairway plot. The solid lines denote the median estimate of the effective population size when the parameter r is 0.5, 0.6, and 0.7. The translucence bands indicate 95% confidence intervals.

**FIGURE 6.**
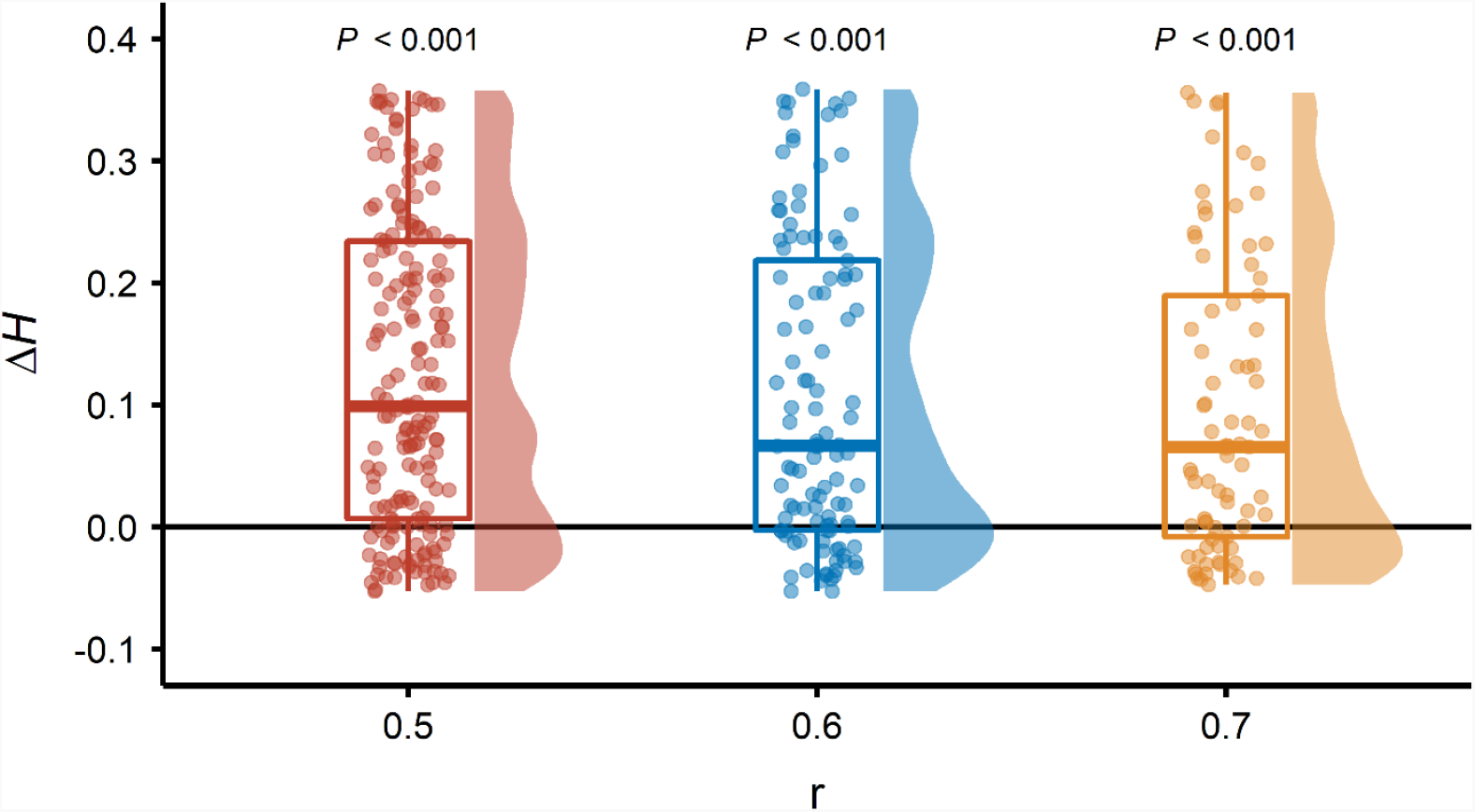
Heterozygosity-excess (*ΔH* = *H*_E_ - *H*_Eq_) based on INEST. The values of *ΔH* in all SNPs with the setting of parameter r = 0.5, 0.6, and 0.7 are represented by dots. The significances (*P* values) in heterozygote excess (*ΔH* > 0) based on Wilcoxon signed-rank test are denoted.

## Discussion

In this study, we revealed 1) the extremely low level of genetic diversity of *G. yakushimensis*, 2) the gentle genetic differentiation across the Yakushima population despite the patchy distribution, and 3) the small population size due to the recent bottleneck and no dramatic increase in population size over the long term.

We demonstrated that *G. yakushimensis* had extremely low levels of genetic diversity and a high level of inbreeding coefficient compared to those of its congener (Table 1). Surprisingly, the genetic diversity across only nine individuals from a single population of *G. triflora* exceeded that across 353 individuals of *G. yakushimensis*. This result was consistent with that of previous studies that showed that relatively rare species have lower genetic diversity than that of common species (Broadhurst et al., 2017; Chung et al., 2018; Gibson et al., 2008; Hamrick & Godt, 1996; Nybom & Bartish, 2000). The limited gene flow across a narrow range (details in the next paragraph) might resulted in inbreeding among closely related individuals. Because inbreeding can lead to the reduction in reproductive fitness (Losdat et al., 2014; O’Grady et al., 2006; Reed & Frankham, 2003), the negative impacts on population dynamics of *G. yakushimensis* related to inbreeding depression is a concern in the near future. The decrease in patch size and connectivity between patches might result in further excellence in inbreeding.

There was less genetic differentiation among the fragmented patches of *G. yakushimensis* across Yakushima Island. IBD was significant only across the narrow range (up to 140–160 m), and the constant pattern of pairwise genetic distance was detected over this range, suggesting that gene flow occurs across the 140–160 m range. The granitic rocks, the habitats of *G. yakushimensis*, are densely distributed in the center of Yakushima Island. Thus, there is gene flow between adjacent habitats, and the patchy habitat network of *G. yakushimensis* might be maintained. These results indicate that *G. yakushimensis* on Yakushima Island comprises almost a genetically single population. *Gentiana yakushimensis* was mainly pollinated by bees (H. Abe, personal observation). The foraging range of bees was at least 100 m for species with the smallest body size (about 1 mm of intertegular span) (Greenleaf, Williams, Winfree, & Kremen, 2007). Generally, bees prioritize the foraging of neighboring flower patches after foraging within a patch (Woodgate, Makinson, Lim, Reynolds, & Chittka, 2017). Thus, the pollination behavior by bees might gently connect patchy habitats and maintain a low level of genetic differentiation.

The population of *G. yakushimensis* was likely to experience the range expansion during or after the LGM and the recent reduction of effective population size according to the demographic inferences and the bottleneck tests (Figure 5 and figure 6). This result was consistent with the previous studies which demonstrated the range expansion of alpine plants during or after the LGM (Ikeda et al., 2020; Ikeda, Yakubov, Barkalov, & Setoguchi, 2018; Schönswetter et al., 2005). Although these previous studies showed the range expansion into the areas that were glaciated during the LGM, Yakushima Island was not glaciated during the LGM (Tsukada, 1985). Based on the results of the present study, it was not possible to distinguish whether the increase in population size was due to a decrease in the vegetation and forest limit associated with the cold climate during the LGM and/or an expansion into the area with the highest altitude related to the warming after the LGM. The low level of genetic diversity in *G. yakushimensis* was assumed to be due to the significant impact of the recent bottleneck. There are three possible reasons for this result: illegal digging (Japanese Ministry of Environment, 2020), grazing by sika deer (Takatsuki, 1990, 2009; Tsujino & Yumoto, 2004), and a habitat area reduction by the uplift of forest and vegetation lines by warming (Japanese Ministry of the Environment, 2017). The high intensity of grazing by mammalian herbivores negatively affected the maintenance of population in the congener of *G. yakushimensis* (Moschetti, Besnard, Couturier, & Fonderflick, 2020). The high density of the sika deer in Yakushima Island (Tsujino & Yumoto, 2004) might lead to the decrease in population size. In Yakushima Island, because the forest cover and density of *C. japonica* increased in the high-altitude area from 1963 to 2014 (Japanese Ministry of the Environment, 2017), it is inferred that the increase in the vegetation limit of shrubs and dwarf bamboos reduces the habitat area of *G. yakushimensis*. Because the seedling establishments in the *Gentiana* species can be limited by the high level of light competition (Ekrtová & Košnar, 2012; Miller, Geddes, & Mardon, 1998), the uplift of vegetation cover is one of the factors strongly influencing the population size via competitions. In addition, because the maximum population size of *G. yakushimensis* estimated to be approximately 9,000 (Figure 5), one of the possible reasons for low genetic diversity is the long-term maintenance of a small population size. While the recent bottleneck was supported by the demographic inferences, the reduction in population size based on the estimated median was low level (Figure 5), which might be attributed to the long lifespan of the genus *Gentiana* (more than 15 years) (Oostermeijer, Veer, et al., 1994). The longevity generally causes a time lag between reducing the apparent population size and the decline in genetic diversity (Fuller & Doyle, 2018; Reisch et al., 2017). This time lag might lead to a further reduction in the genetic diversity of *G. yakushimensis* in the near future.

*G. yakushimensis* has an extraordinarily low level of genetic diversity, resulting in vulnerability to environmental changes and human disturbances (Reusch et al., 2005; Willi et al., 2006). Moreover, the effective range of gene flow in this species is extremely narrow. In species with limited dispersal ability, a high level of patch connectivity can enhance gene flow, decreasing the extinction probability (Dornier & Cheptou, 2012). In the congener of *G. yakushimensis*, the low level of genetic diversity resulted in the reduction in offspring fitness (Oostermeijer, van Eijck, & den Nijs, 1994). Thus, the loss of small patches may threaten the maintenance of fitness in the entire population. Therefore, we emphasize the conservation of large and small patches of *G. yakushimensis*. In the present study, we provided information on the genetic diversity and population demography of *G. yakushimensis*. The accumulation of the genetic information of a lot of endangered species can be useful for the conservation prioritization (Garner et al., 2020), such as the conservation project of plant species with extremely small populations in China (Ma et al., 2013; Ren et al., 2012). For an outlook, we require the assessment of population stability by long-term monitoring for the changes in genetic diversity and population size to identify the factors influencing the reduction in genetic diversity and the length of the time lag between the decrease in apparent population size and effective population size, since a drastic reduction in effective population size can occur in the near future.

## Acknowledgments

The authors are grateful to Yuji Isagi, the project leader of the Environment Research and Technology Development Fund. The authors would like to thank Kenshi Tetsuka, Arata Yoichi, Toshihiro Saito, Tatsuko Tetsuka, and Kengo Fuse for sampling plant materials. We also thank our laboratory colleagues for helping with the analysis and an earlier draft of this manuscript, especially Kenji Seiwa, Yu Fukasawa, Kimiyo Matsukura, Takuya Konuma, and Rimpei Nagaoka. This research was supported by the Environment Research and Technology Development Fund (F-091, 4-1605 and 4-1902) of the Environmental Restoration and Conservation Agency of Japan. Permission to conduct research in the study area was provided by Kagoshima Prefecture, the Japanese Ministry of Environment, the Japanese Ministry of Education, Culture, Sports, Science and Technology, and the Japanese Forestry Agency.

## Author’s Contributions

N.I.I. performed the molecular experiments and analyses and led the writing of the manuscript; Y.T., A.M., and H.A. contributed to the molecular experiments and analyses; S.K.H. and Y.S. contributed to the analyses and writing; and N.I.I., H.A., and Y.S. designed the research.

---

**FIGURE S1.**
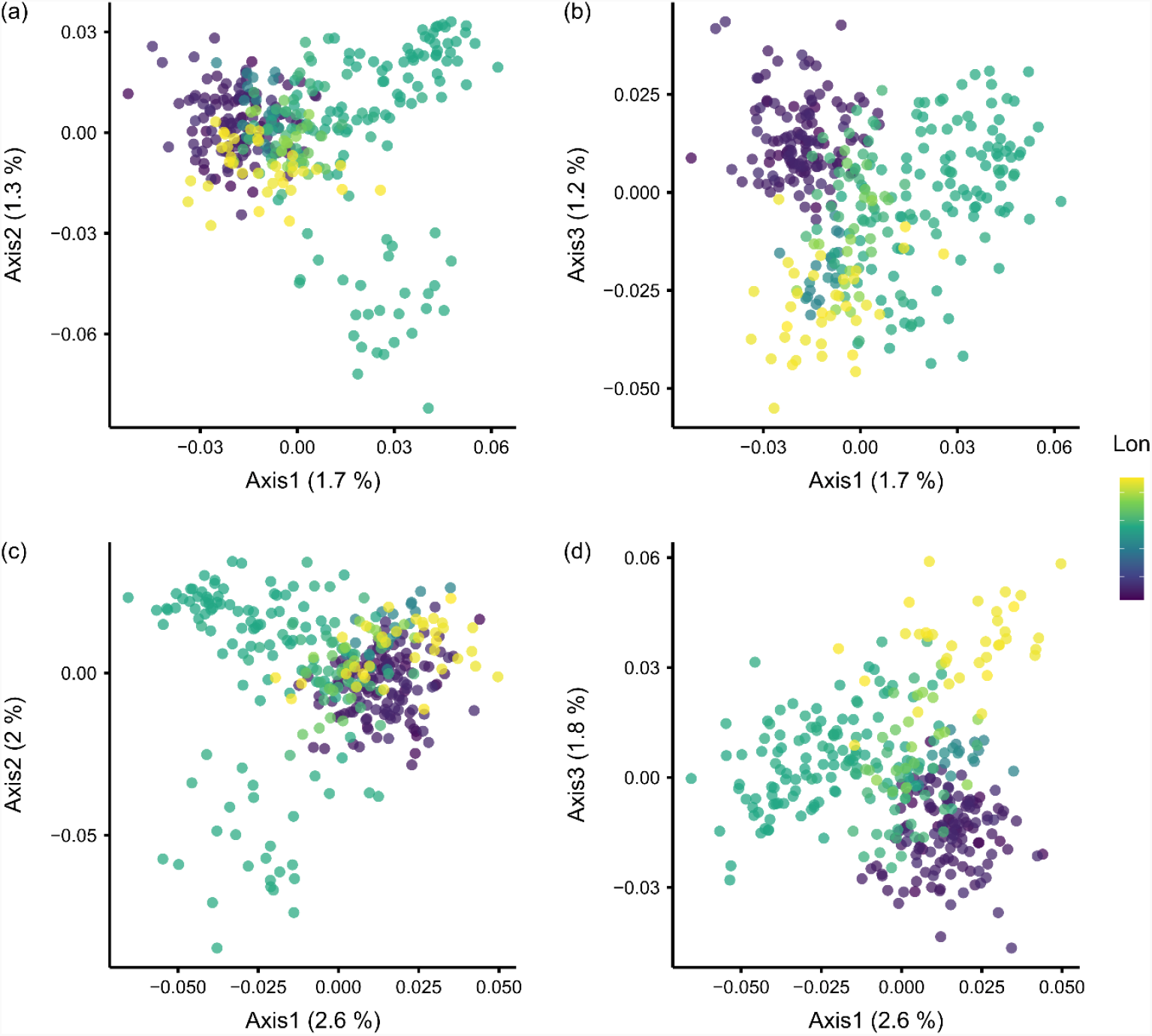
Principal coordinate analysis (PCoA) score plots based on individual-based genetic distance of *Gentiana yakushimensis* when the parameter r is 0.6 (a, b) and 0.7 (c, d). (a, c), axis 1-2; (b, d), axis 1-3. Colors denote the location (longitude) of each individual.

